# Vesicle-bound regulatory RNAs are associated with tissue aging

**DOI:** 10.1101/2021.05.07.443093

**Authors:** Fabian Kern, Thomas Kuhn, Nicole Ludwig, Martin Simon, Laura Gröger, Natalie Fabis, Abdulrahman Salhab, Tobias Fehlmann, Oliver Hahn, Annika Engel, Marcus Koch, Jana Koehler, Katarzyna Winek, Hermona Soreq, Gregor Fuhrmann, Tony Wyss-Coray, Eckart Meese, Matthias W. Laschke, Andreas Keller

**Author notes:** These authors have equally contributed to the study and are listed in lexicographic order (Fabian Kern, Thomas Kuhn, Nicole Ludwig, Martin Simon). These authors contributed equally as senior authors to the study (Matthias W. Laschke, Andreas Keller). Correspondence should be addressed to Andreas Keller.

## Abstract

Previous work on murine models and human demonstrated global as well as tissue-specific molecular aging trajectories in solid tissues and body fluids^1–8^. Extracellular vesicles like exosomes play a crucial role in communication and information exchange in between such systemic factors and solid tissues^9,10^. We sequenced freely circulating and vesicle-bound small regulatory RNAs in mice at five time points across the average life span from 2 to 18 months. Intriguingly, each small RNA class exhibits unique aging patterns, which showed differential signatures between vesicle-bound and freely circulating molecules. In particular, tRNA fragments showed overall highest correlation with aging which also matched well between sample types, facilitating age prediction with non-negative matrix factorization (86% accuracy). Interestingly, rRNAs exhibited inverse correlation trajectories between vesicles and plasma while vesicle-bound microRNAs (miRNAs) were exceptionally strong associated with aging. Affected miRNAs regulate the inflammatory response and transcriptional processes, and adipose tissues show considerable effects in associated gene regulatory modules. Finally, nanoparticle tracking and electron microscopy suggest a shift from overall many small to fewer but larger vesicles in aged plasma, potentially contributing to systemic aging trajectories and affecting the molecular aging of organs.

## Introduction

Understanding and controlling molecular hallmarks of age-related processes in higher organisms promises to greatly improve the quality of life^1^. For humans, aging is frequently studied using easily accessible biospecimens such as blood, serum or urine. Consequently, the scientific community generated models for a broad spectrum of molecular physiological and pathophysiological processes from different molecule types. For example, studies rely on long-lived individuals^2^, serum proteomic profiling^3^, small RNA patterns in blood cells^4,11^, or the exploration of epigenetic control of aging clocks^5^. Likewise, deeper profiles such as gene expression fingerprints are available for different tissues^6^. Murine models facilitate the analysis of such processes thanks to their restricted influence of a heterogenous genetic background and varying lifestyles as compared to humans. Thus, organism-wide RNA-sequencing data of major organs and cell types across the mouse lifespan are available as unique resource to study aging^7,8^. The available data suggest complex aging patterns, including both linear and non-linear effects that are either specific for organs or follow more global organism-wide trajectories. These observations indicate a systemic and well-orchestrated exchange of information and molecules between organs. Specifically, extracellular vesicles (EVs) such as exosomes are postulated to play an important role^9,10^, e.g., hypothalamic stem cells seem to control aging through exosomal miRNAs^12^. Recently, targeted intervention of exosomal transfer of miRNAs from osteoclast to chondrocytes was described as promising method to slow or even inhibit osteoarthritis in mice^13^, as also summarized in a news and views article ^14^. Further, several studies address the relation of EVs with aging in a systematic manner^15–18^. Also of special interest are studies investigating the change of vesicle bound non-coding RNAs depending on (treatment) interventions such as caloric restriction^19^. However, these studies are often limited in their analysis scope by considering only one small RNA class at a time and even restricting themselves to a subset of well characterized representatives. Moreover, an inherent restriction often is the limited sample count, frequently leading to pooling of biosamples and blurring fine-grained signals.

These and other issues complicate the analysis of EVs and their molecular load. Especially in the context of EVs in cancer, common pitfalls in purification have been summarized by Schekman and co-workers^20^ with the correct nomenclature of EVs, purification and other aspects elucidated in great detail. Similar limitations as described in the context of cancer in this review also apply for other diseases, matrices and organisms.

For advancing our understanding on how systemic non-coding RNAs are associated with aging, we sought to discern differences between the molecular information included in EV cargo and freely circulating non-coding RNAs. We thereby balance the tradeoff of having sufficient material for high-throughput sequencing of individual plasma samples at the highest purity of input material. For each individual mouse, plasma and EV samples we sequenced non-coding RNAs and contrasted them by computational approaches.

## Results

To uncover age-related dynamic processes and to model the information exchange involved we sequenced both free circulating non-coding RNAs and vesicle-bound non-coding RNAs from individual mice. The molecular profiles are available at five time points across the average lifespan between two to 18 months in two to four replicates per age group and biospecimen type (***Fig. 1A, Supplemental Table 1***). For the plasma and the EV RNA samples, we sequenced on average 38 million reads and mapped them to ten different non-coding RNA classes. In total, the feature set includes 80,688 different non-coding RNAs in mice, mostly piRNAs, circRNAs, lincRNAs and miRNAs (***Fig. 1B***). The very first aspect encompassed the distribution of molecules from the different RNA classes. While tRNA fragments were highly represented both in exosomes and as freely circulating molecules, piRNAs showed sharply lower levels in both specimen types (***Fig. 1C***). However, dominantly for circRNAs and rRNAs significantly different amounts as freely circulating molecules compared to vesicle-bound RNAs were observed. Notably, this analysis has a quantitative and RNA class centric view but does not yet consider whether the representative within the classes match across sample types. For example, only a small number of piRNAs are expressed both in plasma and exosomes even though general abundance was high. Considering the sample type overlap for each class, indeed the most significant difference was observed between circulating and vesicle-bound piRNAs (***Fig. 1D***). Similarly, we report large differences in the content of RNA molecules from snRNAs, snoRNAs, and scaRNAs. In contrast, detected tRNA fragments, lincRNAs, rRNAs and circRNAs are often shared between the two specimen types. In sum, our data thus argue for a type-specific expression patterns that differ significantly between non-coding RNA classes in a quantitative and qualitative manner.

**Figure 1:**
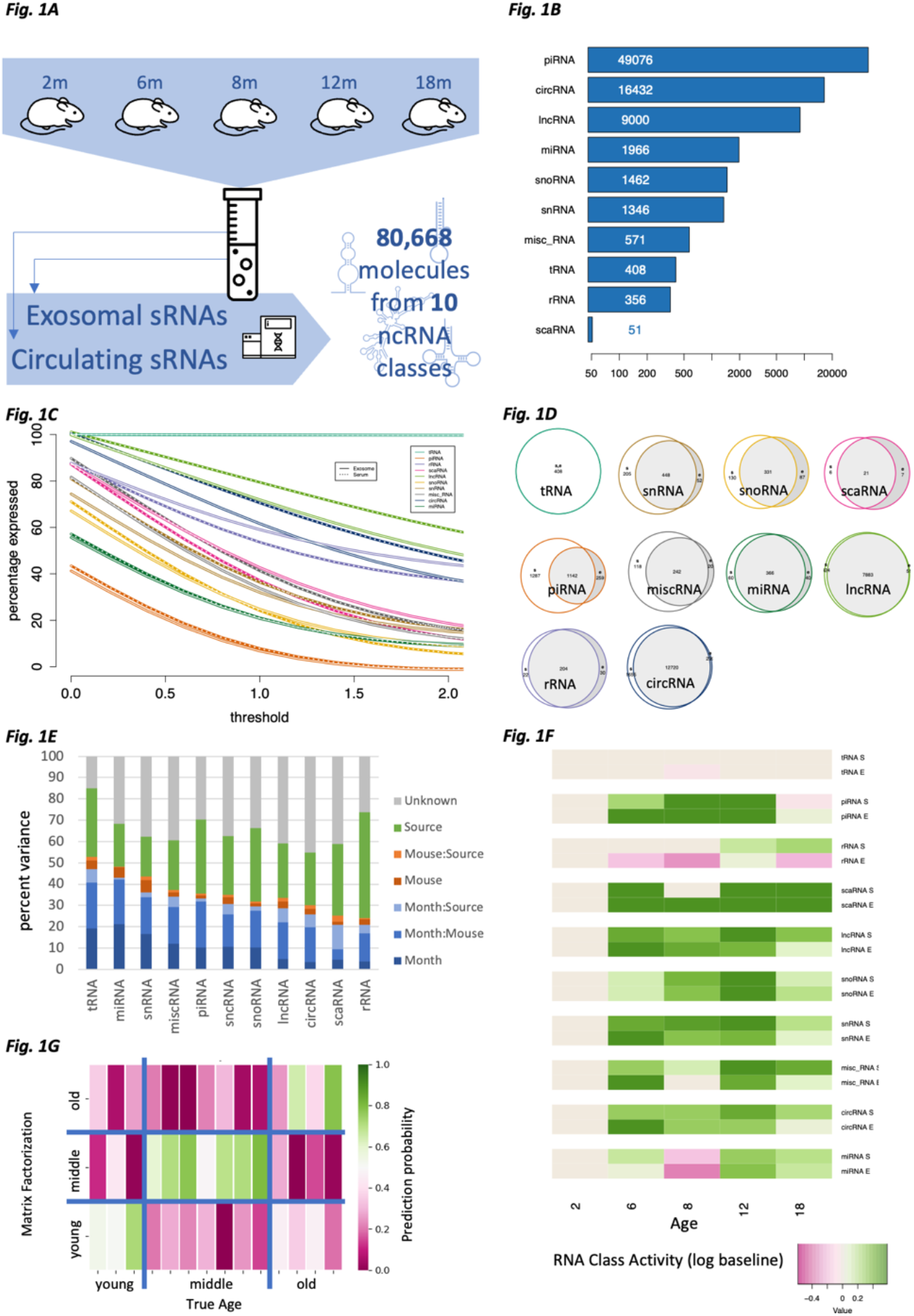
Distribution of sequencing reads into their mapped non-coding RNA classes and their general relation to aging. (**A**) Study set-up. We profiled vesicle and plasma samples from mice in 5 age groups and sequenced 80,668 non-coding RNAs from 10 classes. (**B**) Overall distribution of molecules to the 10 non-coding RNA classes. (**C**) Fraction of representatives per RNA class dependent on an expression threshold. Dashed lines are serum, solid lines vesicle samples. (**D**) Overlap of expressed RNAs in plasma and vesicles as area proportional Venn diagrams. (**E**) percent of total data variance attributed to different parameters such as the age (month), individual mice or specimen type (source). Columns are sorted according to decreasing fraction of variance attributed to the age. (**F**) Relative expression of the different RNA classes per timepoint and sample type as compared to the baseline (2 months). Green means higher expression and purple lower expression. The upper row per RNA class is serum, the lower row is vesicles. (**G**) Prediction of age by non-negative matrix factorization. The color code represents the probability (trust) in the prediction, x-axis represents the true age group, y-axis the predictions.

Our initial hypothesis was to test for unique aging trajectories within and between non-coding RNA classes wrapped in vesicles or freely circulating in plasma. Thus, we computed the proportions of variance in the RNA counts that can be explained by available sample metadata, i.e., either by age of the mice, the specimen type and donor mice identity, or linear combinations of such. Here, tRNA fragments and miRNAs clearly stood out in terms of fraction of variance explained by the age (***Fig. 1E***). In comparison, the lowest information content with respect to age was obtained by scaRNAs and rRNAs. Importantly, the individuality factor of each donor mice used for this study was comparably small, independent of the RNA class. To uncover the actual relationship between each RNA class and mice age, we set the expression at month 2 as baseline and modelled whether activity increases or decreases over time in EVs and plasma, separately. As one result, we found remarkably similar dynamics of change for vesicles and free circulating RNAs.

The largest overall age-related differences exist in rRNAs where the overall amount increases for free circulating molecules with aging but the vesicle loading of rRNAs decreases. This substantial difference in the specimen types also explains the high proportion of variance attributed to the sample type annotation (rightmost green bar in ***Fig. 1E***). Our data further indicate a strong aging signal in EVs and in plasma, with varying strengths, again depending on the RNA class (***Fig. 1F***). As our previous analyses emphasized the role of tRNA fragments, we investigated the expression profiles in an unbiased manner and performed a classification into three age groups (young, 2 months; middle aged, 6-8 months; old, 12-18 months). We modelled this classification task as optimization problem through non-negative matrix factorization, computing probabilities for each sample to belong to each of the three age groups. We then assigned each sample to the age group with highest probability. For both plasma and micro-vesicles, we got varying prediction accuracies, once again with the best results for tRNA fragments with a remarkable accuracy of 86% (***Fig. 1G***).

Taken together, our analyses suggest non-coding RNAs to exhibit specific age trajectories, both in qualitative and quantitative aspects. Moreover, the data pinpoint at substantial differences in case of circulating RNAs and vesicle bound RNAs where the best correlation with age was observed for tRNA fragments. This immediately poses the question whether loading of vesicles follows biologically relevant environmental mechanisms. To discover such potential patterns, we next performed a fine-granular and molecule-centric analysis.

From the 80,668 unique non-coding RNA molecules in *Mus musculus* included in our analysis, 23,052 (28.6%) are stably expressed in plasma and vesicles (***Supplemental Table 2***). We then computed the linear Pearson correlation as well as the non-linear distance correlation for each of the investigated RNAs.

Based on this information, we estimated for each RNA whether it is linearly correlated with age, nonlinearly correlated with age or not correlated with age at all for plasma and vesicles separately. For both sample types, freely circulating in plasma (***Fig. 2A***) and vesicle bound (***Fig. 2B***), the linear component was dominant and online few exceptions with non-linear trajectories occurred. The amplitude and frequency of non-linear RNAs were both slightly enriched in EVs. Interestingly, we also observed a slight enrichment towards negative correlation in EVs. Both aspects argue for a more characteristic abundance of RNA levels in aging EVs as compared serum. Having observed non-coding RNAs that are positively and negatively correlated with age in vesicles or freely circulating further called for exploring whether the up- and down-regulated molecules show similar compositions in the two specimen types. In total, 27% and 22% are increasing and decreasing with age in EVs and serum, respectively, slightly differing from what would be expected by a random distribution (cf. methods). Intriguingly, however 39% of the 23,052 expressed non-coding RNAs are negatively correlated in EVs with age but positively correlated in plasma and with 12% behaving vice-versa (***Fig. 2C, Supplemental Table 2***). To seek common patterns for the increasing and decreasing expression of non-coding RNAs we clustered the expression in EVs and in plasma separately and extracted RNA clusters from the dendrogram. For each cluster we then computed the average linear and non-linear correlation with aging and finally calculated the overlap of the sample types. Our analysis confirmed the strong and orchestrated decrease of correlation with age in EVs as compared to plasma **(*Fig. 2D*)**. The vesicle clusters are enriched in the lower left corner, indicating the significant trend towards negative correlation with age in EVs. Further, the data reveals a substantial age-related loss in linear correlation as compared to non-linear correlation. To validate the origin of these signals we inspected all concordant and discordant non-coding RNAs and provide specific examples for markers clearly increasing with age in plasma and vesicles (*miR-466i-5p, **Fig. 2E***), decreasing with age in plasma and vesicles (*Gm16701, **Fig. 2F***), increasing with age in plasma but decreasing with age in vesicles (*Gm20756, **Fig. 2G***), and lastly decreasing with age in plasma but increasing in vesicles (*miR-690, **Fig. 2H***). We further examined whether the patterns hold for all the 10 non-coding RNA classes or rather if they are class specific. Here, the specificity of patterns for the different non-coding RNA classes was astonishing. For example, 94% of tRNA fragments increased with age in both plasma and in EVs. Also, 54% of rRNAs decreased with age in plasma but increased with age if vesicle-bound. Inversely, 42% of circRNAs increased with age in plasma but increased with age if vesicle-bound. Also, other RNA classes revealed distribution patterns significantly deviating from the 25% per group as expected by chance. For example, 82% of miRNAs increased with age in EVs **(*Fig. 2I*)**.

**Figure 2:**
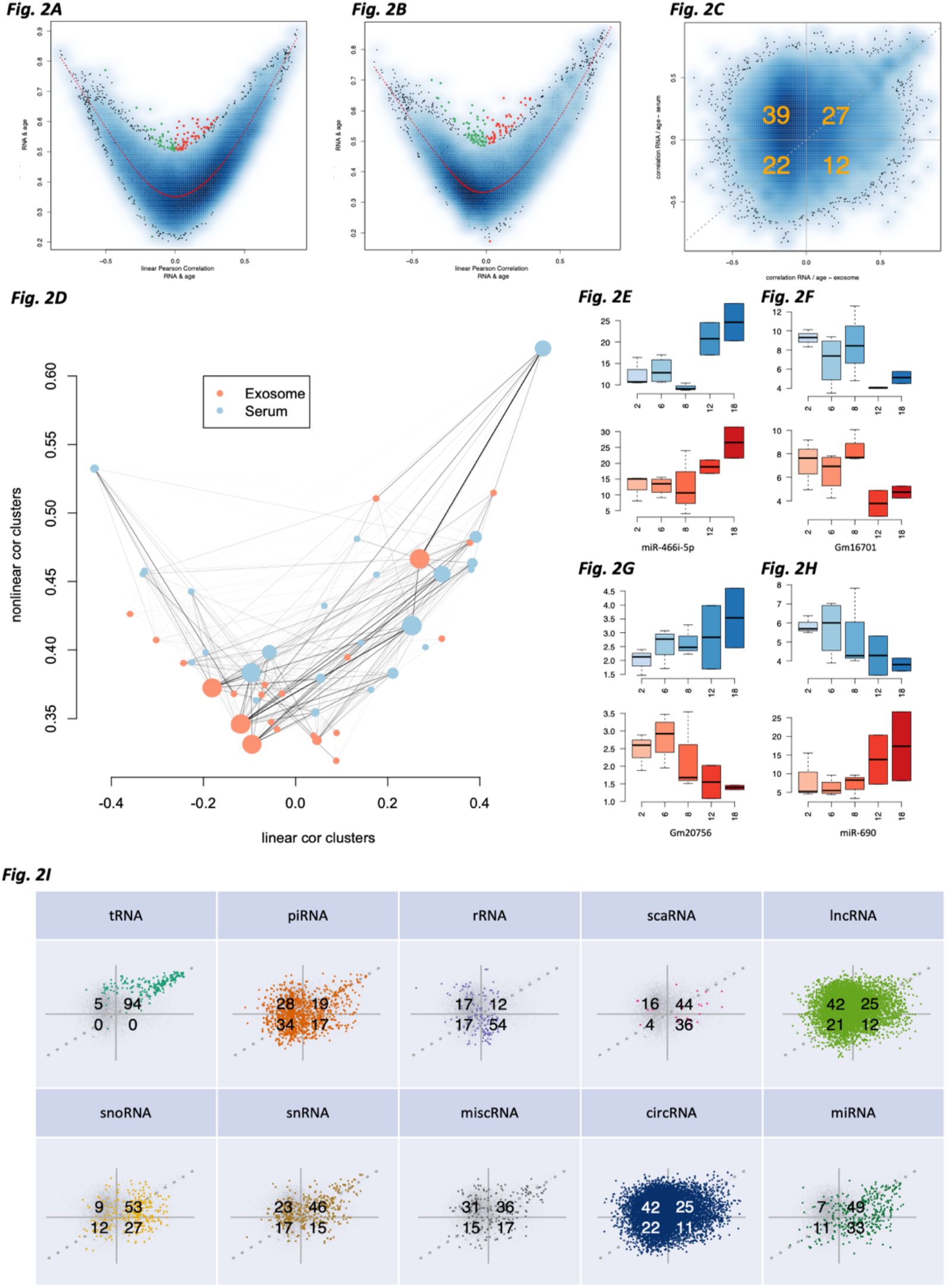
Correlation of non-coding RNAs with aging in EVs and in serum. (**A**) For each non-coding RNA, the x-axis represents the Pearson correlation and the y-axis the distance correlation with age in serum. The red line is a smoothed spline, and the colored dots (green, negatively; red, positively) are correlated with age in a predominantly non-linear manner. (**B**) The same information as in panel (A) but for EVs. (**C**) Scatter plot showing the Pearson correlation in vesicles (x-axis) in relation to the Pearson correlation in plasma (y-axis). Orange numbers represent the percentage of points in each of the four quarters. The data suggest a shift to negative correlation with age in vesicles. (**D**) Non-coding RNAs in plasma and EVs were clustered and resulting clusters were attributed with the average linear and non-linear correlation. Solid lines represent a match of non-coding RNAs, the thicker the line the more non-coding RNAs match between a plasma and vesicle cluster. The diameter of the points represents the cluster size. Most EV clusters accumulate in the lower left corner. (**E-H**) Examples of non-coding RNAs that increase or decrease with age. The x-axis represents the age, the y-axis the expression of the selected non-coding RNA (orange, EV; blue serum). (**I**) Confusion matrix scatter plots (see also panel (A) split by the RNA classes.

In the light of the regulatory mechanism of miRNAs typically repressing gene expression^21^ and further knowing that mRNA levels tend to decrease with age, we next asked whether the miRNAs increasing or decreasing with age in vesicles or plasma exhibit a distinct function in a pathway-specific manner. For each gene ontology category^22^ we computed an enrichment score for the miRNAs in EVs and in serum. Next, we used the list of miRNAs sorted by their correlation with age to perform cutoff-free miRNA set enrichment analysis^23,24^. A direct comparison provided strong evidence supporting the notion that miRNAs correlated with age in vesicles are significantly enriched in biochemical categories as compared to those in plasma **(*Fig. 3A*)**. This finding is in line with our initial hypothesis that EVs are specifically loaded with non-coding RNAs that exert biological processes in remote sensing cells. To understand the nature of these processes, we compared the 16 categories that are at least three orders of magnitude more significant in vesicles as compared to plasma to the two being at least three orders of magnitude more significant in serum. In the former, the strongest enrichment was found for the protein heterodimerization activity, neural crest cell migration, negative regulation of inflammatory response, receptor internalization, positive regulation of neuroblast proliferation, the mitochondrial envelope, the positive regulation of DNA-templated transcription and the TORC2 complex (all with adjusted p-value < 5×10^−5^ in vesicles and p-value > 0.01 in serum). The categories with higher significance in plasma are cellular response to BMP stimulus (p=65×10^−5^ vs. 0.11) and negative regulation of myotube differentiation (p=65×10^−7^). Distinct pathways being specifically enriched in vesicles opens the question on potential effects in gene regulation in different tissues.

**Figure 3:**
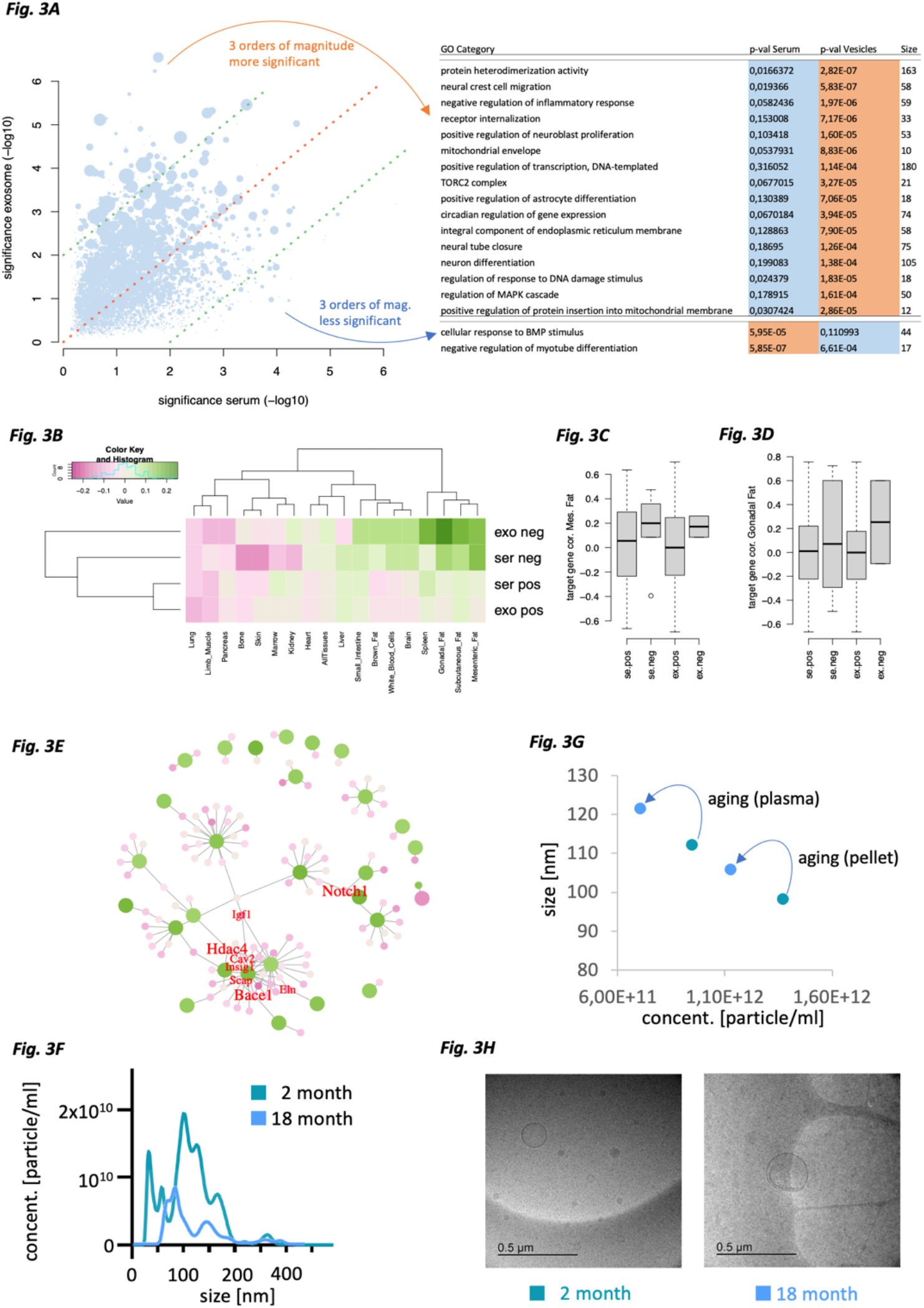
Pathway results and nanoparticle tracking. (**A**) miRNA pathway enrichment for age-related miRNAs in plasma (x-axis) and vesicles (y-axis). Each dot is one pathway, the size represents the number of miRNAs associated with the pathway. The red dashed line is the bisector, the green lines mean two orders of magnitude higher significance in plasma and vesicles, respectively. The pathways with at least three orders of magnitude difference are listed on the right. (**B**) For miRNAs that are positively and negatively correlated with age in plasma and vesicles, the correlation with age of target genes from Tabula Muris Senis across 17 tissues is shown. Expected is that miRNAs going down with age have target genes going up with age and vice versa. (**C,D**) For two fat tissues the target gene correlation with age is detailed for the four groups shown as rows in (B). (**E**) Target network. Large green dots depict genes, small pink dots represent miRNAs and lines delineate regulatory events between miRNAs and genes. The color shading represents the correlation with age and the hub genes targeted by at least three miRNAs are annotated in red. Relative font sizes represent the number of miRNAs targeting the respective gene. (**F**) Size histogram of two selected examples from the nanoparticle tracking analysis, one young and one old mouse. (**G**) Average concentration and size of nanoparticles in young and old mice before and after ultracentrifugation. For both types of analysis we observe the shift towards fewer but larger vesicles in aged mice. (**H**) Selected electron microscopy images of one young and one old mouse plasma sample show EVs and a limited background signal.

The core hypothesis of our work stipulates specific loading of EVs with non-coding RNAs, first and foremost miRNAs, enabling the performance of specific function and gene regulation in remote cells. Previously, we established a bulk- and single cell murine tissue aging atlas, Tabula Muris Senis^7,8^. In these studies, we report both linear and nonlinear aging trajectories in gene expression signals. Similar to the findings on non-coding RNAs observed here, the associated genes cluster with coherent biological functions (among others, extracellular matrix regulation, unfolded protein binding, mitochondrial function, and inflammatory and immune response). The expression patterns are consistent across tissues, differing only in the amplitude and age of onset. Especially fat tissues showed early aging signals of biochemical pathways similar to those observed in the miRNA pathway analysis described above. It was previously shown that miRNAs target genes in a pathway specific manner^25–27^. Thus, for miRNAs associated with age we extracted validated target genes from miRTarbase^28^ and evaluated the correlation of potential targets with age annotations from Tabula Muris Senis (***Fig. 3B***). In this context, the expected pattern is a negative correlation of target genes where miRNAs increase with age and vice versa, and as such was confirmed for several organs, including mesenteric fat, gonadal fat, the brain’s white blood cells and brown fat. Likewise, the tissue independent aging patterns matched our expectations. While fat tissues generally showed the best concordance, other tissues such as the lung or pancreas did not. The target gene correlation for mesenteric and gonadal fat verified the increased correlation with age for miRNAs decreasing with age and vice versa (***Fig. 3C/3D***). Intriguingly, this effect was more pronounced for gonadal fat, however only for EV bound miRNAs and less so for serum. Translating the miRNAs and genes to a proper target gene network identified 8 core genes: *Notch1*, *Bace1*, *Hdac4, Igf1, Eln, Cav2, Insig1*, and *Scap (**Fig. 3E***), possibly reflecting physiological relevance for both signaling networks and epigenetic processes.

In sum, our analysis reinforces the central role of fat in early aging processes, indicating that vesicle bound miRNAs have important contributions. Having demonstrated that the cargo of EVs seems to be biologically relevant further opens the question whether it is just the cargo of the EVs or whether the nature of vesicles is altered.

We finally carried out nanoparticle tracking analysis (NTA) and Cryogenic electron microscopy (cryo-EM) on a new cohort of mice. Thereby we focused on the extreme ages, i.e., studied four mice of the youngest and four mice of the oldest age group. A general shift in the distribution from more and smaller EVs present in young animals to fewer but larger EVs in old animals was observed (***Fig. 3F, Supplemental Fig. 1***). We performed this analysis for samples with and without ultracentrifugation to minimize noise in the data. Although this analysis is challenging and apoptotic bodies might blur the image, we find for both sample types with and without centrifugation identical patterns: aging leads to a decrease of the particle concentration and at the same time to an increase of the average particle diameter (***Fig. 3G***). Both the differential loading and the different size and number of EVs seem to add to the altered molecular aging profiles across tissues. Although only in a qualitative manner, also cryo-EM supported the thesis of different sizes and counts for vesicles in young and old plasma (examples are presented in ***Fig. 3H***, all available cryo-EM images are shown in **Supplemental Fig. 2**).

## Discussion

While our study presents intriguing new insights into the correlation of EVs and the molecular loading of EVs with non-coding RNAs in the context of aging, it is important to mention possible limitations and how we took them into account. First, the purification of EVs is challenging and bears many pitfalls^20^. Generally, the more purification steps are applied the less material is left, requiring a pooling of samples. We decided to have the maximal purity possible while still leaving sufficient material for individual sequencing of small RNAs, avoiding any pooling. As stated by Schekman et al. ^20^, healthy skepticism concerning the possible connection between exosomal miRNAs and control of gene expression in target cells will remain until functional cell culture and animal studies are conducted with exosomes purified by rigorous and quantitatively documented procedures. While NTA and cryo-EM do not replace such purification, they demonstrate a limited background noise in the used samples. A second limitation comes down to the molecular measurement and annotation of the molecules. Having reached a high sequencing depth from low input volumes, the data are mapped to the standard reference databases. Whether read molecules, e.g., mapping to piRNAs actually represent functional piRNAs and not fragments or reads mapping to miRNAs annotated in miRbase are representing functional miRNAs, is only partially known, calling for further functional validation experiments. These factors have to be addressed in targeted studies with larger cohorts of mice and using orthogonal molecular profiling technologies such as RT-qPCR.

## Online Methods

### Animals

For the non-coding RNA sequencing experiments, we used female C57BL/6N mice at the age in months of 2 (n = 4; body weight (bw): 19-20 g), 6 (n = 4; bw: 25-29 g), 8 (n = 4; bw: 23-26 g), 12 (n = 2; bw: 31 g) and 18 (n = 4; bw: 34-42 g). To assess the spread of vesicles in fat tissues, a second cohort of female C57BL/6N mice was used with an age of 2 (n = 4; bw: 19-20 g) and 18 months (n = 4; bw: 29-41 g). The animals were housed in groups on wood chips as bedding in the conventional animal facility of the Institute for Clinical & Experimental Surgery (Saarland University, Homburg/Saar, Germany). They had free access to tap water and standard pellet food (Altromin, Lage, Germany) and were maintained under a controlled 12-h day/night cycle. This animal study was approved by the local State Office for Health and Consumer Protection and conducted in accordance with the Directive 2010/63/EU and the NIH Guidelines for the Care and Use of Laboratory Animals (NIH Publication #85-23 Rev. 1985).

### Blood sampling

The mice were anesthetized by an intraperitoneal injection of ketamine (100 mg/kg bw; Ursotamin®; Serumwerke Bernburg, Bernburg, Germany) and xylazine (12 mg/kg bw; Rompun®; Bayer, Leverkusen, Germany). Subsequently, they were fixed on a heating pad in supine position. After midline laparotomy, a maximal volume of blood (~700-1000 μL) was taken from the vena cava and transferred into plasma tubes (Sarstedt, Nümbrecht, Germany). The blood samples were then centrifuged at 20°C and 10.000 xg for 5 min and the resulting plasma was stored at −80°C until further use.

### Purification of EVs

200 μl of mouse plasma were transferred to 1 mL open-top thickwall polypropylene ultracentrifugation tube (Beckman-Coulter, USA) and diluted with 800 μL of phosphate-buffered saline to prevent the tube from collapsing in the ultracentrifuge vacuum. Samples were centrifuged for 2 h, at 4°C at 100,000 x g using the Type 50.4 Ti fixed-angle rotor (Beckmann-Coulter, USA). Supernatants were carefully removed, and the pellets were resuspended in 20 μL of phosphate-buffered saline. Samples were stored at −80°C until further analyses.

### RNA extraction

Total exosomal RNAs were isolated semi-automated using the miRNeasy Micro kit (Qiagen, Hilden, Germany) and Qiacube isolation robot according to the manufacturer’s recommendations with addition of 2 μL RNase-free glycogen (20 mg/mL, Invitrogen, Carlsbad, CA, USA) to facilitate RNA precipitation. RNA concentration was measured using Qubit™ microRNA Assay Kit (ThermoFisherScientific, Waltham, MA, USA).

### High-throughput RNA-sequencing

Isolated RNA samples were analyzed by Agilent small RNA Chips and 2 ng each (exosomes and supernatants) were used for Illumina compatible library preparation using the D-Plex small RNA Kit (Diagenode, BE). The kits employ 3’-poly A tailing and template switch-based cDNA generation using UMI (unique molecular identifier) tagged template switch oligos. After PCR amplification involving 13 cycles, libraries were purified from TBE-PAGEs. Illumina sequencing was carried out on a HiSeq2500 platform using the High Output mode for 96 cycles.

### Nanoparticle-tracking analysis

1 μl of serum was diluted in 1199 μL, 1 μL of the resuspended pellet was diluted in 999 μL of phosphate-buffered saline, in order to achieve a final concentration between 20 and 120 particles/frame. Samples were then measured on NanoSight (Malvern, UK) at a camera level of 15. For each sample, three captures of 30 s were acquired. Videos were then analyzed at a detection threshold of 5 using NTA 3.4 software.

### Cryo-Transmission Electron Microscopy

3 μL of each sample were transferred to a holey carbon film-coated copper grid (Plano S147-4), blotted for 2 s, and plunged into undercooled liquid ethane at –165 °C (Gatan Cryoplunge3). The grid was then, under liquid nitrogen, transferred to a cryo-TEM sample holder (Gatan model 914). Low-dose bright-field images were acquired at –170 °C, using a JEOL JEM-2100 LaB6 Transmission Electron Microscope and a Gatan Orius SC1000 CCD camera.

### Bioinformatics

The sample primary processing was performed with miRMaster^29^ using standard parameters. As output, miRMaster generated a list with the expression of 80,668 RNAs from 10 RNA classes. The data were normalized to expression in one million reads and further processed with R (R 4.0.4 GUI 1.74 Catalina build (7936)). The Venn diagrams were generated using the *eulerr* package from R. Mapping the fraction of variance to different parameters was done using the principal variance component analysis (*pvca*) package. Splines were computed using the *smooth.spline* function with seven degrees of freedom. Color palettes were generated using the *RColorBrewer* package. Smoothed scatter plots were computed using the *smoothScatter* function setting the point number to 500. Clustering was done for highest expressed non-coding RNAs (at least 5 reads per million in at least one sample) using the scaled expression matrix (z-score of each feature). The clustering was performed with the *hclust* function, using the Euclidean distance measure. Clusters were extracted by cutting the dendrograms at 1/1.25 of the maximal height. Heatmaps of target genes were computed using the *heatmap.2* function. The network visualization has been performed using *iGraph*. As input for the network analysis, targets from miRTarbase ^28^ were used, however, restricted to strong evidence targets. To compute the statistical concordance of RNAs correlated with aging across sample types, a random background distribution with respect to positive and negative correlation was assumed. Briefly, a random distribution would mean that close to 25% of non-coding RNAs is consistently positively regulated in serum and EVs, 25% is consistently negatively correlated with age and 25% in each are positively correlated in the one and negatively correlated in the other specimen type.

### Matrix factorization

We predicted the age of samples with respect to three age groups “young” (2 months), “middle” (6-8 months) and “old” (12-18 months). To this end, the expression patterns were split in 20 individual matrices, for each of the 10 non-coding RNA classes and for plasma and EVs. We first normalized the given non-negative Matrix D by dividing all elements by the maximum value in *D*. To obtain the probabilistic regarding the age groups, we decomposed the matrix *D* into two further matrices *T* and *P*, where *P* gives us the desired probabilities. *T* stands for the matrix of the typical age group vectors, i.e., in each entry of a column there is a value representing all entries at this position of all samples belonging to this age group. The matrix *P* contains the probabilities of each sample to each age group respective to their typical vector in *T*. We formulated the non-negative matrix factorization as the optimization problem:

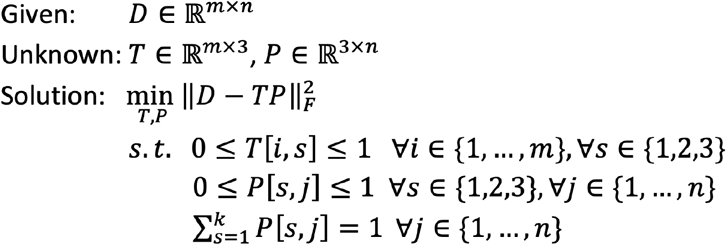

The first two constraints require all entries of the matrices *T* and *P* to lie between 0 and 1. Since we were interested in the percentage of a sample belonging to the three age groups, we also required all columns of the matrix P to sum up to 1 using a numerical solver^30^.

We then classified each sample by choosing the index with the highest entry of each column in P and assigned the index as a label to each one. However, the rows of P were invariant to permutations. Here, this means that it is not clear which label corresponds to which age group. Furthermore, a measure of quality for the results was needed. Using the known age, we could construct a ground truth for each sample and calculated the classification accuracy for every permutation. The ground truth used was “young” corresponding to two-month-old mice, “middle” to six- and eight-months-old and “old” to twelve- and 18-month-old. Finally, we chose the permutation labels which maximize the accuracy and obtain a measure of quality.

## Data availability

All sequencing data are freely available from the Sequence Read Archive SRA [reviewer link to the data is available upon request].

## Code Availability

The primary data analysis has been performed using miRMaster. miRMaster is available as web service. The source code of miRMaster is not publicly available because of licensing issues. The other analyses have been carried out using standard R packages that are freely available (see Methods Section).

## Author contributions

FK contributed to data analysis, data interpretation, and writing the manuscript. TK performed the vesicle extraction experiments. NL contributed to the study set up and the interpretation of the analysis. MS contributed to the sequencing of the samples and to the study set up. LG contributed to the versicle purification experiments. NF contributed to the sequencing experiments. AS contributed to the sequencing experiments. TF contributed to the analysis of primary sequencing data. OH contributed to the analysis of the tissue experiments from TMS. AE performed the matrix factorization analysis. MK performed the Cryo EM analysis. JK contributed to the matrix factorization analysis and the interpretation of the results. KW and HM contributed to the tRNA fragments analysis and interpretation. GF contributed to the vesicle extraction and to the study set up. TWC contributed to the analysis and interpretation of the data in the context of the tissue experiments from TMS. EM contributed to the study set up and the interpretation of the results. MWL performed the mouse experiments. AK contributed to study set up, data interpretation and manuscript writing.

## Acknowledgements

We appreciate the support of the mouse facility at Saarland university. We appreciate input from the members of the Keller lab.

## Funding

This study has partially been funded by the Regional Government of the Saarland and by Saarland University.

Supplementary Materials are available online

