## Supplemental Figures 1&2 for "Vesicle-bound regulatory RNAs are associated with tissue aging"

*Supplemental Figure 1*

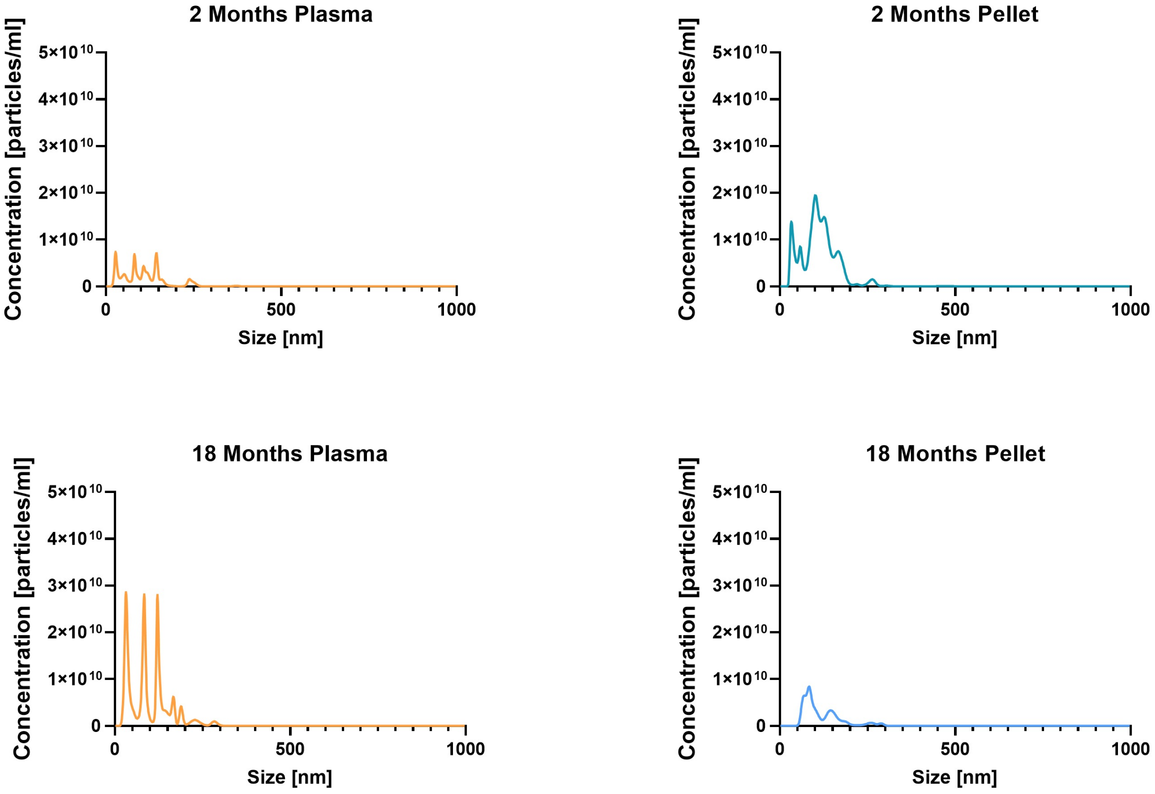


NTA analysis for 2 months plasma and pellet compared to 18 months plasma and pellet.

*Supplemental Figure 2*

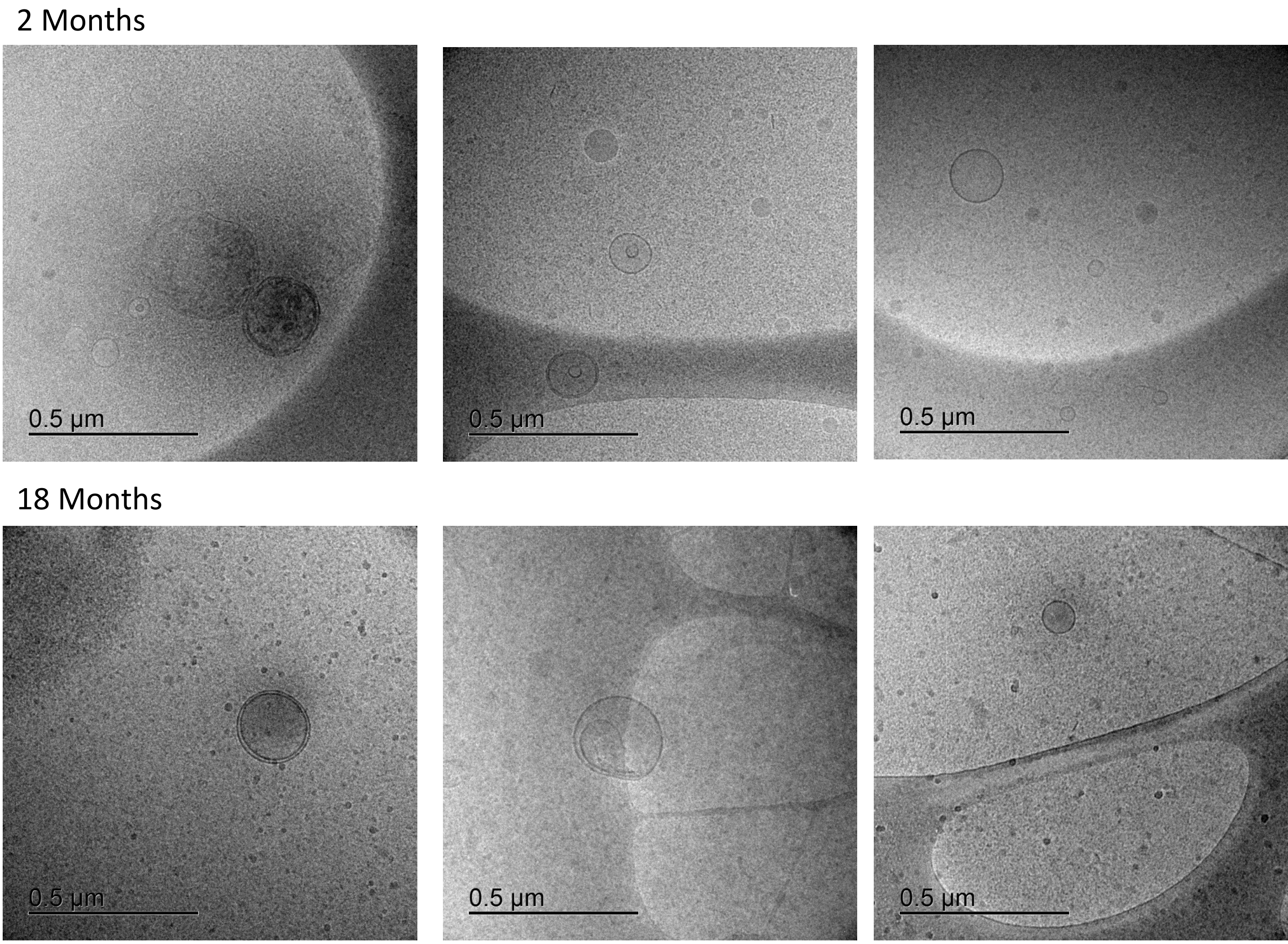


Cryo EM Images of three 2-motnh and three 18 month samples
